# Cytokinin promotes jasmonic acid accumulation in the control of maize leaf growth

**DOI:** 10.1101/760405

**Authors:** Aimee N. Uyehara, Angel R. Del Valle-Echevarria, Charles T. Hunter, Hilde Nelissen, Kirin Demuynck, James F. Cahill, Georg Jander, Michael G. Muszynski

## Abstract

Growth of plant organs results from the combined activity of cell division and cell expansion. The coordination of these two processes depends on the interplay between multiple hormones that determine final organ size. Using the semidominant *Hairy Sheath Frayed1* (*Hsf1*) maize mutant, that hypersignals the perception of cytokinin (CK), we show that CK can reduce leaf size and growth rate by decreasing cell division. Linked to CK hypersignaling, the *Hsf1* mutant has increased jasmonic acid (JA) content, a hormone that can inhibit cell division. Treatment of wild type seedlings with exogenous JA reduces maize leaf size and growth rate, while JA deficient maize mutants have increased leaf size and growth rate. Expression analysis revealed increased transcript accumulation of several JA pathway genes in the *Hsf1* leaf growth zone. A transient treatment of growing wild type maize shoots with exogenous CK also induced JA pathway gene expression, although this effect was blocked by co-treatment with cycloheximide. Together our results suggest that CK can promote JA accumulation possibly through increased expression of specific JA pathway genes.

**One sentence summary:** Cytokinin-signaling upregulates the jasmonate biosynthesis pathway, resulting in jasmonate accumulation and influences on maize leaf growth.

## INTRODUCTION

Growing plants accumulate biomass over time through the integration of cell division and cell expansion. These processes produce biomass by increasing cell number (cell division) and increasing final cell volume (expansion). In eudicot leaves, the placement and timing of cell division, expansion, and differentiation determines the pattern of leaf growth. In many model plants, leaf growth follows a basipetal pattern where differentiation starts at the distal tip of the leaf and finishes near the proximal base (Gupta and Nath, 2016; Conklin et al., 2019). Other growth patterns include acropetal growth (differentiation starts at the proximal base), diffuse growth (differentiation occurs evenly across the leaf without respect to cellular proximal-distal position), and bidirectional growth (differentiation begins at both the distal tip and proximal base) (Gupta and Nath, 2016).

Growth is controlled, in part, by signaling between plant hormones. Plant hormones are molecular messengers with low molecular weights that regulate growth, development, and defense (Santner and Estelle, 2009; Wolters and Jürgens, 2009; Frébort et al., 2011; Huot et al., 2014). Generally, plant hormones can be divided into two classes: growth hormones and defense hormones. Classical growth hormones include, cytokinin (CK), gibberellins (GA), brassinosteroids (BR) and auxin (Huot et al., 2014). These hormones have been ascribed functions in cell proliferation, stem elongation, seed germination, and organ elongation respectively. Classical defense hormones include salicylic acid (SA), jasmonic acid (JA), and ethylene (ET), and are responsible for the majority of signaling in response to pests and pathogens (Huot et al., 2014). Coordination between growth and defense pathways is necessary for appropriate allocation of resources in response to environmental stimuli, and is mediated by hormone crosstalk. One described example of crosstalk is the signaling between GA and JA. In the presence of JA, the GA repressor DELLA is released to bind and degrade GA leading to suppression of GA-mediated growth by JA (Hou et al., 2013). In contrast, BR seems to relieve JA-induced growth suppression, suggesting an antagonistic relationship between BR and JA (Huot et al., 2014). Crosstalk has also been shown to occur between SA and auxin. SA represses auxin-mediated growth by repressing the transcription of the F-box protein TIR1/AFB, leading to the stabilization of the auxin repressor AUX/IAA (Wang et al., 2007; Huot et al., 2014). As predicted by the growth-defense tradeoff model, signaling by defense hormones to growth hormones often leads to growth suppression.

Growth of the maize leaf occurs at its base within zones of cell division and expansion that are spatially distinct. The maize leaf contains multiple growth associated hormones which crosstalk to regulate and define the leaf growth zones (Nelissen et al., 2012). The maize leaf develops from the maize leaf initials on the shoot apical meristem. The leaf tip is formed first and is propelled distally by proliferative divisions at the leaf base (Kiesselbach, 1999). In monocot leaves, the basipetal growth mechanism sets up regions of division, elongation, and maturation that are linearly organized and spatially separated into distinct growth zones (Nelissen et al., 2016). The linear organization of the growth zones makes it straightforward to use kinematic analysis to measure the relative contribution of division and expansion to final leaf size (Nelissen et al., 2013). Kinematic analysis provides insight into the complex molecular interactions underlying leaf growth as different hormones have measurable and distinct impacts on the growth zones. This was demonstrated through kinematic analysis of GA biosynthesis mutants in maize (Nelissen et al., 2012). It was shown that increased bioactive GA increases the size of the division zone and determine the spatial location of the division-elongation transition zone (Nelissen et al., 2012). These data also implicated other growth hormones such as cytokinin, auxin, and brassinosteroids as possible players in determining the size of the division zone (Nelissen et al., 2012).

Cytokinin (CK) is a growth promoting hormone that regulates processes such as shoot growth, apical dominance, senescence, and promotion of cell proliferation (Miller et al., 1955; Werner et al., 2001). In dicots, cytokinin promotes leaf growth by stimulating cell division. This has been demonstrated through exogenous CK treatment, overexpression of CK catabolic enzymes, or knockout of CK receptors. For example, decreasing endogenous CK concentration through the overexpression of the CK catabolic enzyme, CYTOKININ OXIDASE (CKX) in *Nicotiana tabacum* reduced leaf size by reducing cell number (Werner et al., 2001). In *Arabidopsis thaliana*, reduction of CK signaling through the knockout of the CK receptor *Arabidopsis* HISTIDINE KINASE 2 (AHK2), AHK3, and CRE1/AHK4 resulted in plants with severely reduced rosette size and a reduced number of cells per leaf (Riefler et al., 2006). Reduced cell number as a result of reduced cytokinin perception or signaling resulted in growth compensation through cell expansion (Werner et al., 2001; Riefler et al., 2006). In contrast, constitutively active CK receptor mutants in *A. thaliana* exhibited larger leaves with more epidermal cells due to either an extended period of mitotic activity, increased mitotic rate, or both (Bartrina et al., 2017).

The role of CK in regulating monocot leaf growth is less clear. In contrast to *Arabidopsis*, maize has seven CHASE-domain histidine kinase receptors (Lomin et al., 2011; Steklov et al., 2013). The maize mutant *Hairy Sheath Frayed1* (*Hsf1*) is the only CK receptor gain-of-function monocot mutant. *Hsf1* is a semidominant mutant with an EMS-induced mutation in the cytokinin receptor, *Zea mays HISTIDINE KINASE1 (ZmHK1)*, an orthologue of *AtHK4 (Bertrand-Garcia and Freeling, 1991; Muszynski et al., 2019)*. Characterization of *Hairy Sheath Frayed1* (*Hsf1*) demonstrated the role of increased CK signaling had on leaf patterning, leaf size, and epidermal cell fate (Bertrand-Garcia and Freeling, 1991; Muszynski et al., 2019). Although CK typically promotes cell division and growth, increased signaling (hypersignaling) of CK in *Hsf1* mutants reduced leaf growth compared to wild-type siblings (Muszynski et al., 2019). The effect of reduced CK on monocot growth was indirectly observed through transgenic overexpression of zeatin O-glucosylzeatin, an enzyme that inactivates and sequesters CK through the addition of a sugar moiety (Pineda Rodo et al., 2008). Homozygous *Ubi:ZOG1* maize lines showed CK deficiency phenotypes such as reduced growth and interestingly, a feminized tassel (Pineda Rodo et al., 2008).

Jasmonic acid (JA) is an established plant growth regulator involved in processes such as leaf senescence, plant defense, and male fertility (Yan et al., 2014). Linolenate lipoxygenase (LOX) catalyzes the first step of JA biosynthesis from chloroplast membrane phospholipids (Lyons et al., 2013). The resulting hydroperoxy octadecadienoic acids are further converted into (+)-7-iso-JA via allene oxide synthase (AOS), allene oxide cyclase (AOC), 12-oxophytodienoic reductase (OPR) and three cycles of ß-oxidation (Lyons et al., 2013). Bioactive JA-Ile is formed through the conjugation of an amino acid by the jasmonate amido synthetase (JAR) (Lyons et al., 2013). Catabolism of JA-Ile occurs through the oxidation of JA-Ile by the cytochrome CYP94B enzyme (Lunde et al., 2019). Research on JA’s role as both a defense and plant growth regulator is aided by biosynthesis and signaling mutants. In maize, mutants for LOX, OPR, and CYP94B include *tasselseed1, opr7-5 opr8-2*, and *Ts5* respectively (Acosta et al., 2009; Yan et al., 2012; Lunde et al., 2019). These mutants add to a growing body of research that establishes JA as a growth repressor. Initial studies showed that exogenous JA application to rice seedlings reduced seedling leaf size (Yamane et al., 1980). More recently, wound induction of JA and analysis of *Arabidopsis* JA biosynthesis mutants have shown that JA suppresses cell proliferation leading to reduced leaf size with fewer and smaller epidermal cells (Zhang and Turner, 2008; Noir et al., 2013).

Here, we show that CK signaling reduces cell division in the leaf growth zone through promotion of JA accumulation. To do this, we used exogenous hormone treatments, hormone biosynthesis and signaling mutants, kinematic analysis of leaf growth, and expression analysis. Altogether, our data identified a previously unrecognized connection between cytokinin and the defense hormone JA in regulating maize leaf growth.

## RESULTS

### *Hsf1* mutants have a reduced growth phenotype

We have previously shown that *Hsf1*/+ mutants have smaller leaves and that exogenous CK treatment can phenocopy this effect (Muszynski et al., 2019) (Figure 1A, Supplemental Figure S1). To further characterize this reduced growth phenotype, leaf size and growth rate and duration of seedling leaf #4 was determined for *Hsf1*/+ and wild type sibling plants in three different genetic backgrounds (Figure 1A and B, Supplemental Figure S1). In all three backgrounds, *Hsf1*/+ leaf #4 blade length was reduced 10-20% compared to their wild type siblings (Figure 1A, Supplemental Figure S1A). The B73 background was used for the remainder of the studies. Consistent with a reduced blade size, leaf elongation rate (LER) was also reduced by 20-25% across the three backgrounds (Figure 1B, Supplemental Figure S1B and C). Interestingly, leaf elongation duration (LED) was slightly increased for *Hsf1*/+ leaf #4 which may account for the fact that reduction in leaf size is not as great as the reduction in LER would predict. To determine the cellular basis underlying this growth rate reduction, kinematic analysis was performed on *Hsf1*/+ and wild type siblings in the B73 genetic background (Nelissen et al., 2013). Kinematic analysis showed that *Hsf1*/+ mutants had fewer dividing cells in the division zone and thus had a smaller division zone in leaf #4 compared to wild type (Figure 1C). These data suggested that CK hypersignaling in *Hsf1*/+ mutants reduced cell divisions in the leaf growth zone, which slowed growth rate, resulting in a smaller leaf.

**Figure 1.**
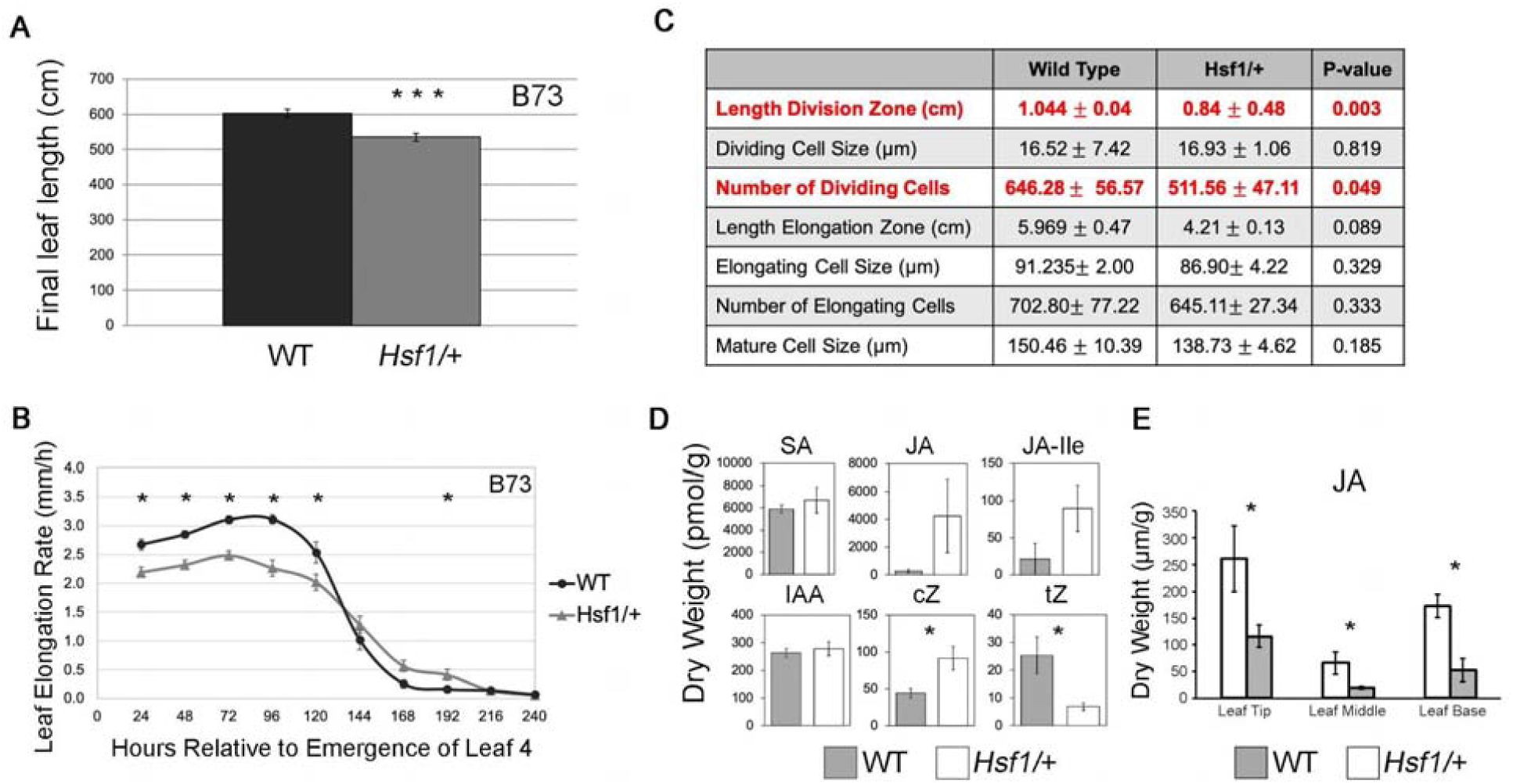
*Hsf1* growth and phytohormone phenotypes. **(A)** Barplots of WT and *Hsf1*/+ final leaf lengths. Error bars = SE. **(B)** Average leaf elongation rate (LER) of leaf #4 of *Hsf1*/+ and WT-siblings in the B73 inbred background. Asterisks mark significant difference P < 0.05. Error bars = SE. **(C)** Kinematic analysis comparing growth zones of the *Hsf1*/+ mutant and its WT-sibling. **(D)** Two-week old whole-seedling hormone profile of *Hsf1*/+ and WT-siblings. SA, Salicylic Acid; JA, Jasmonic Acid; JA-Ile, Jasmonic Acid Isoleucine; IAA, Indole-3-Acetic Acid; cZ, *cis*-Zeatin; tZ, *trans*-Zeatin. **(E)** Jasmonic Acid (JA) concentration across leaf nine at steady-state growth. The leaf was divided into three sections (leaf base, leaf middle, and leaf tip). Leaf base includes the growth zone. White columns are *Hsf1*/+ and gray columns are WT-sibling.

### *Hsf1*/+ accumulates jasmonic acid in growing maize leaves

Plant hormones are known to exert their function through crosstalk with other hormones (Santner and Estelle, 2009; De Vleesschauwer et al., 2014; Huot et al., 2014). To determine if CK hypersignaling in the *Hsf1* mutant was affecting other hormones that may impact growth, differences in phytohormone content were determined by high performance liquid chromatography of WT and *Hsf1* whole seedlings. *Hsf1*/+ accumulated 4-16-fold more of JA-Ile and JA respectively compared to wild type (Figure 1D). A few other hormones showed modest differential accumulation but not in a pattern consistent with the *Hsf1* reduced growth phenotype. To obtain a better spatial resolution of the elevated JA content in *Hsf1*, mature leaf blade #9 was sampled, divided into thirds along the proximal-distal axis, and JA content determined. Consistent with the whole seedling data, JA content was elevated 2-3-fold across the entire *Hsf1*/+ leaf (Figure 1E). CK had not previously been shown to affect JA content but JA was known to inhibit cell division in eudicots, and thus provided a possible mechanism by which the *Hsf1* mutation conditioned reduced growth (Yamane et al., 1980; Zhang and Turner, 2008; Noir et al., 2013). This prompted us to assess the effects of JA on maize leaf growth.

### JA pathway genes are upregulated in the leaf growth zone of *Hsf1* mutants

Given that JA content was increased in *Hsf1* mutants, we assessed whether the expression of some JA pathway genes were increased in the *Hsf1* leaf growth zone. The growth zone of leaf #4 at steady-state growth was partitioned into 5 mm subsections providing a high-resolution spatial sampling through the division zone, first transition zone and elongation zone (Figure 2). Subsections were collected in triplicate and transcript levels for select JA pathway genes were measured by quantitative real-time PCR. These genes were chosen to broadly survey key steps in JA biosynthesis and because mutants are available for some (Gao et al., 2008; Yan et al., 2014) (Figure 2). We found the JA biosynthetic genes *ts1, ZmAOC2* and *ZmOPR7* were significantly upregulated in the division zone of *Hsf1/+* (Figure 2). This suggested that increased JA accumulation was due to increased expression of at least one JA biosynthetic gene(s) in the division zone of *Hsf1* mutant leaves. In addition, the JA-responsive gene *ZmMYC2* had higher expression throughout the entire growth zone in *Hsf1/+*, suggesting increased JA levels were being perceived by the JA signaling pathway (Figure 2). Overall, the expression data supports the hypothesis that CK signaling promotes JA accumulation through increased expression of JA biosynthetic genes. However, we cannot discern the influence JA feedback might have on these results. Since JA accumulation is increased in the growth zone of *Hsf1* leaves and it is known that JA positively regulates its own biosynthesis (Pauwels et al., 2009; Ahmad et al., 2016), we are not able to determine the specific influence CK has on JA pathway gene expression in a “high” JA genotype.

**Figure 2.**
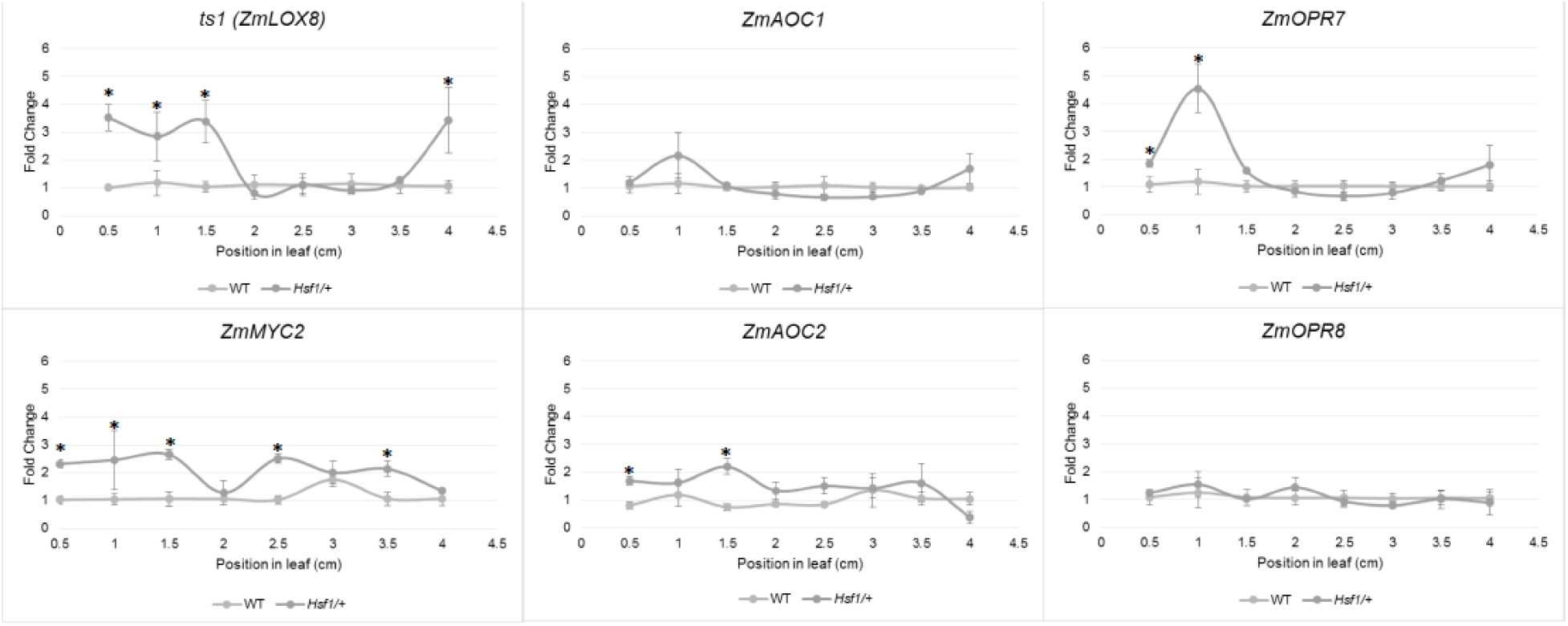
JA pathway genes are up-regulated in the growth zone of *Hsf1* leaves. RT-qPCR of key JA biosynthesis and signaling genes across the division zone in *Hsf1/*+ and wild type leaf #4 at steady state growth.

### Exogenous jasmonic acid treatments reduce leaf growth rate in maize

To test if increased expression of JA biosynthetic genes could be responsible for reduced leaf growth in the *Hsf1* mutant, B73 inbred maize seeds were transiently treated with 1 mM JA and effects on seedling leaf growth were assessed (see Materials & Methods for details). Exogenous JA treatment of germinating maize seeds resulted in a 25-30% reduction in sheath and blade length for seedling leaves #1 to #4 (Figure 3A, Supplemental Figure S2 and Supplemental Table S1). JA treatment also promoted reductions in blade width which varied between 9-20% depending on leaf number (Figure 3A and Supplemental Table S1). Similar to effects of JA in other plant systems, these data indicated that JA treatment can reduce leaf size in maize seedlings.

**Figure 3.**
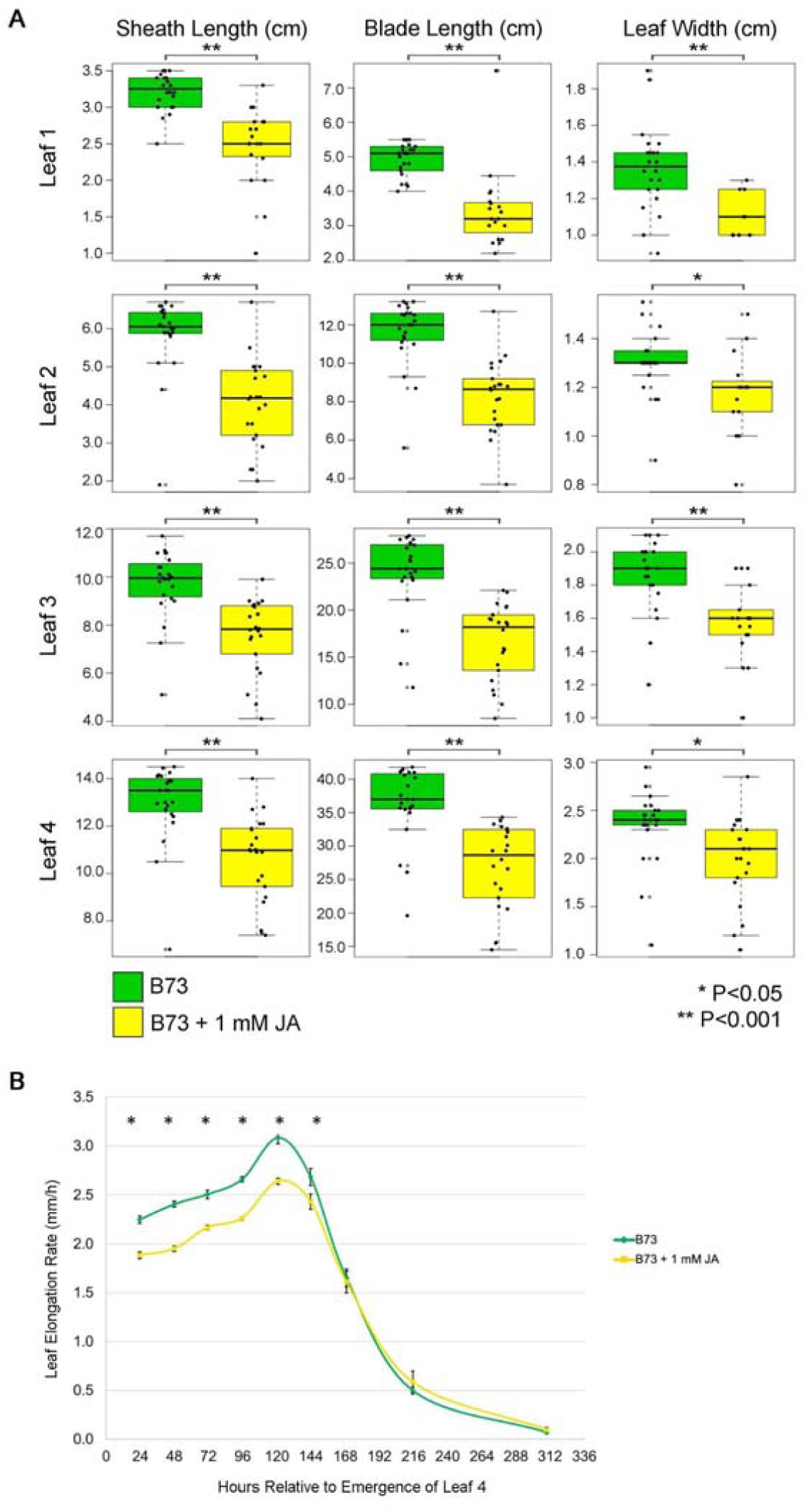
Effect of JA on B73 growth. **(A)** Boxplots of sheath length, blade length, and blade width in control and 1 mM JA treatments of leaf #1-4. Horizontal bars represent the maximum, third quantile, median, first quantile, and minimum values respectively. Each dot is a plant (B73, n=23; B73 + JA, n=22). **(B)** Average leaf elongation rate (LER) of leaf #4 at steady-state growth of seedlings in control compared to 1 mM JA treatment groups. Error bars = SE. Asterisks mark significant differences of LER between treatments at each time point by Student’s t-test p-value ≤ 0.05 (B73, n=27; B73 + JA; n=22).

The JA mediated decrease in leaf size could have resulted from a reduction in growth rate or the duration of growth or both. To distinguish the cause of leaf size reduction, the LER and LED were determined for leaf #4 from B73 seedlings treated with JA, as described above. While both control and JA treated plants maintained steady state growth for five days, JA-treated seedlings had a pronounced reduction in LER compared to control throughout the period of steady-state growth (Figure 3B). No obvious change in LED was observed. To determine the minimum time of JA treatment required to elicit the observed growth reduction, germinating B73 seeds were treated with 1 mM JA for 1, 6, 12, 24 and 48 hours (see Methods for details). Consistent decrease in blade length and width were observed only after 48 hours of JA exposure for leaves #1 to #3 (Supplemental Figure S3 and Supplemental Table S2). Thus, exogenous JA treatment for at least 48 hours could decrease maize leaf size by reducing growth rate. These treatments supported a possible role of JA in reducing *Hsf1* growth rate.

### *Hsf1* is less responsive to exogenous jasmonic acid treatment

Because *Hsf1* mutant leaves have more JA and are smaller than wild type, we hypothesized that leaf size of *Hsf1* mutants would be less responsive to exogenous JA treatment than wild type siblings or the B73 inbred. To test this, we treated germinating seeds that were segregating 50% *Hsf1*/+ and 50% wild type with 1 mM JA using the standard germinating seed hormone assay. The excessive pubescence *Hsf1* phenotype (increased macrohair density on the abaxial sheath) was not affected by exogenous JA treatments and was 100% concordant with previous molecular genotyping (data not shown). Thus it was a reliable and reproducible method to score seedlings as either *Hsf1*/+ or wild type. As expected from previous analysis, leaf size in untreated *Hsf1*/+ was reduced approximately 20% compared to untreated wild type siblings (Figure 4A, Supplemental Table S3). JA treatment reduced leaf size in both wild type and *Hsf1/+* genotypes compared to their respective controls (Figure 4A). However, the response to JA in *Hsf1/+* plants was not as great as in the JA treated wild type plants, as leaf size reduction was dependent on the leaf tissue and parameter measured. JA treatment reduced wild type sheath length, blade length, and blade width about 15-25%, similar to reductions seen in JA treated B73 seed, although blade #4 width was not affected (Figure 4A and Supplemental Table S3). However, only blade length was consistently reduced (17-25%) in JA-treated *Hsf1*/+ plants, with no reduction in sheath length and inconsistent reduction in blade width (Figure 4A and Supplemental Table S3). These results suggest that in the *Hsf1/+* mutant, blade length but not the other leaf growth parameters are responsive to the JA treatment.

**Figure 4.**
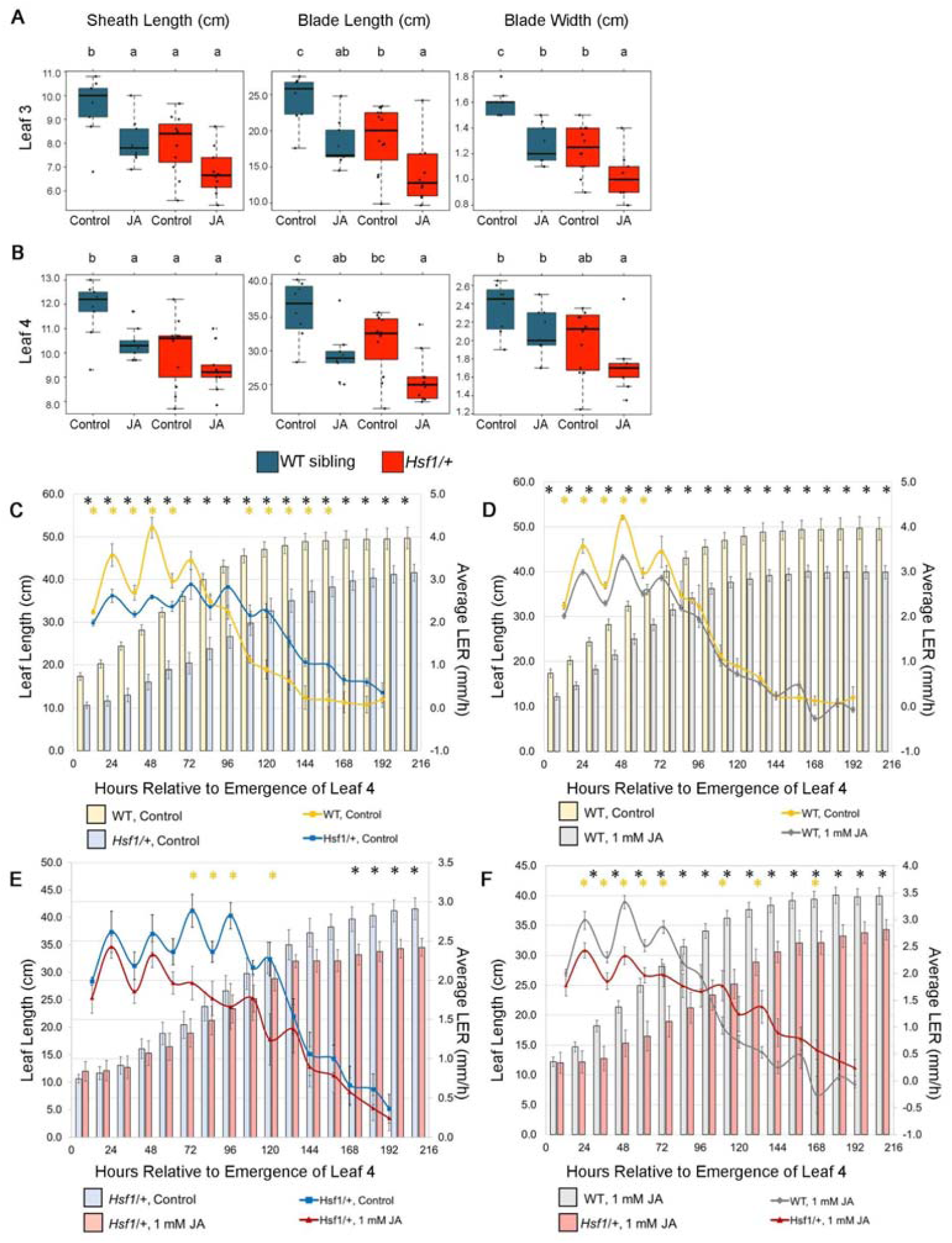
Final leaf size, leaf elongation rate (LER), and leaf elongation duration (LED) of *Hsf1*/+ and WT-siblings treated with 1 mM JA. Boxplots of leaves #3 **(A)** and #4 **(B)** of *Hsf1*/+ and WT-siblings from seedlings grown from germinating seed subjected to a 6-day, 1 mM JA treatment. Horizontal bars represent the maximum, third quantile, median, first quantile, and minimum values respectively. Each dot is a plant (WT Control, n=7; WT JA, n=9; *Hsf1*/+ Control, n=10; *Hsf1*/+ JA, n=9). **(C-E)** LER superimposed over total leaf length. **(C)** LER and leaf lengths of WT and *Hsf1*/+ control treatments. JA treatment comparisons in **(D)** WT, **(E)** *Hsf1*/+, and **(F)** treated *Hsf1*/+ and WT. Significant differences by Student’s t-test are marked by asterisks. Yellow asterisks mark differences in LER and black asterisks mark differences in leaf length. Error bars = SE.

Since JA treatment further reduced *Hsf1/+* blade size, we asked if the JA treatment was affecting growth rate or duration of growth. To do this, LER and LED were determined for leaf #4 of seedlings from 1 mM JA treated 1:1 segregating *Hsf1* and wild type seeds (as above). As seen previously, compared to untreated wild type sibs, untreated *Hsf1*/+ had a reduced LER and extended LED (Figure 4C). Also similar to our results with JA-treated B73, JA-treated wild type LED was not affected but LER was reduced which was especially evident in the first 2.5 days of growth (Figure 4D). In contrast, *Hsf1/+* LER, especially during the first 2.5 days of steady-state growth, was not affected by JA treatment. Instead, LED was reduced by JA treatment in *Hsf1*/+ plants where steady-state growth began to slow starting at 3 days, instead of day 5, and continued to slow until leaf growth stopped by day 8 (Figure 4E). Comparing JA-treated wild type and JA-treated *Hsf1*/+ growth, showed a reduced LER but extended LED for *Hsf1*/+ plants, as was seen for these genotypes without JA treatment (Figure 4F). Thus, although *Hsf1*/+ blade length can be reduced further by JA treatment, it is likely caused by a shortened LED, since LER was not impacted. This can be seen when comparing the actual leaf length (sheath length + blade length) of growing leaf #4 from both genotypes with and without JA treatment (Figures 4C to 4F). Leaf length was reduced at each time point during leaf growth for wild type vs. *Hsf1*/+, for wild type vs. JA-treated wild type, and for JA-treated wild type vs. JA-treated *Hsf1*/+ (Figures 4C, 4D and 4F). In contrast, *Hsf1*/+ vs. JA-treated *Hsf1*/+ showed leaf length was not different until after 7 days of leaf growth, nearly the time growth stopped (Figure 3E). This suggests that in *Hsf1* mutants, where steady-state leaf growth is reduced, possibly by increased JA content, additional JA can only further reduce leaf size by truncating the duration of growth.

### Growth is enhanced in jasmonic acid-deficient mutants

Our data are consistent with previous work showing JA can reduce growth. This implies that reduced endogenous JA accumulation may enhance growth leading to larger leaves. To understand how endogenous concentrations of JA might affect leaf growth and size, we measured leaf size and growth in a number of maize JA deficient mutants (Yan et al., 2012; Lunde et al., 2019). Duplicate genes encode 12-OXO-PHYTODIENOIC ACID REDUCTASE (OPR), a key enzyme in the JA biosynthetic pathway responsible for converting OPDA into (+)-7-iso-JA, which is later modified into bioactive JA (Yan et al., 2012). Plants homozygous for recessive null mutations in both the *opr7* and *opr8* genes are JA deficient, display a feminized tassel or “tasselseed” phenotype, and have longer seedling leaves #1 and #2 (Yan et al. 2012). A single functional *opr* allele at either locus, renders that genotype wild type for JA content and plant phenotype. Using a population that was homozygous null for *opr7* and segregating for a wild type and null *opr8* alleles, we assessed leaf size and leaf growth in JA sufficient and JA deficient genotypes (Figure 5A and 5B). As was shown previously, leaf #1 and #2 sheath and blade lengths of the JA deficient genotype was increased 20%-48%, and leaf #3 and #4 blade length was increased 13%-24% (Figure 5A and B and Supplemental Table S4). Interestingly, sheath length was increased for leaf #3 but decreased in leaf #4 in the *opr7/opr7, opr8/opr8* mutant (Supplemental Table S4). We also noted that blade width increased in leaf #3 and #4 9%-18% in the JA deficient genotype. Overall, in *opr7 opr8* double mutants, increases in sheath and blade length diminished from leaf #1 to #4 but blade width changed from smaller to larger than wild type. Assessment of growth rate in the double *opr7 opr8* mutant revealed an increase in LER and LED compared to the JA-sufficient genotypes (Figure 5B). This suggested the lack of JA increased both the rate and the duration of leaf growth.

**Figure 5.**
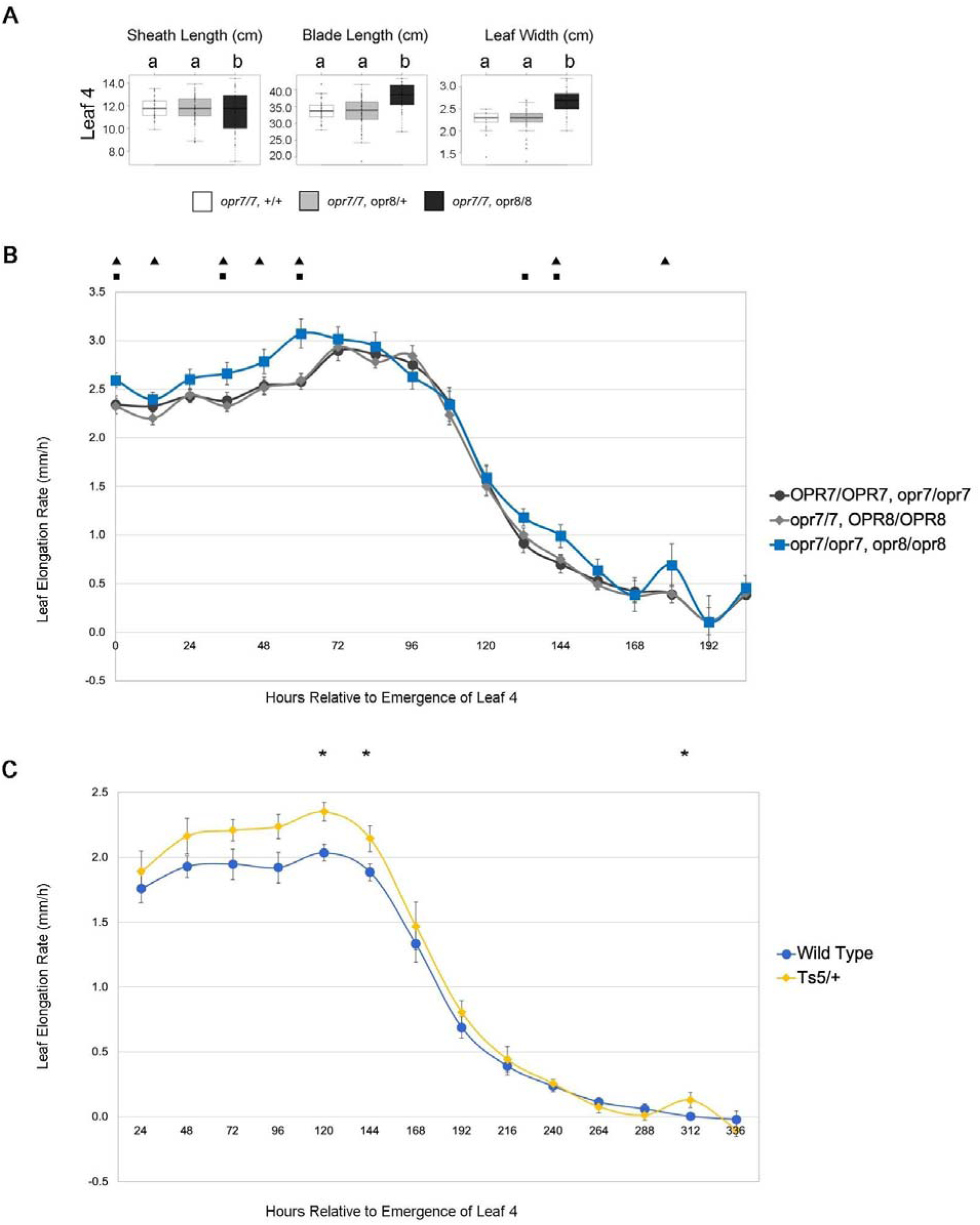
JA deficiency in maize enhances leaf growth. **(A)** Boxplots of sheath length, blade length, and blade width of the JA-deficient *opr7 opr8* double mutant as compared to its JA-sufficient siblings (*opr7/opr7, OPR8/OPR8* and *opr7/opr7, OPR8/opr8*). **(B)** LER of JA-deficient *opr7 opr8* double mutant as compared to its JA-sufficient siblings *opr7/opr7, OPR8/OPR8* (black triangles) and *opr7/opr7, OPR8/opr8* (black squares). Asterisks mark significant difference by Student’s t-test p-value ≤ 0.05. Error bars = SE (*OPR8/OPR8*, n=34; *OPR8/opr8*, n=62; *opr8/opr8*, n=33). **(C)** LER of JA-deficient *Ts5* dominant mutant compared to its JA-sufficient WT-sibling. Asterisks mark significant difference P < 0.05.

To extend the results above, we also measured leaf size and growth in the semi-dominant, gain-of-function *Tasselseed5* (*Ts5*) mutation (Lunde et al., 2019). The *Ts5* locus encodes a cytochrome P450 enzyme, ZmCYP94B1, that oxidizes the bioactive JA-Ile to 12OH-JA-Ile which is less bioactive and *Ts5* mutants express more *ZmCYP94B1* than wild type (Lunde et al., 2019). Thus, *Ts5*/+ plants have a lower JA content than wild type sibs and display the tasselseed phenotype expected for JA deficient mutants. *Ts5/+* was crossed to *Hsf1*/+ and the 1:1:1:1 segregating population was analyzed for LER and LED. LER and LED was measured and plants were genotyped for *Ts5*/+. First we analyzed *Ts5/*+ growth compared to wild type. *Ts5*/+ plants exhibited increased LER compared to wild type and possibly an increase in growth duration (Figure 5C). Consistent with the results from the *opr7 opr8* population, these JA deficient mutants showed increased growth rate, supporting the role of reduced JA promoting leaf growth.

### JA-deficient mutants suppress the reduced leaf growth phenotype in *Hsf1* mutants

Using the population described in Figure 5C, we next compared the LER and LED of single and double mutants. *Hsf1*/+ mutants had reduced LER and an extended LED compared to wild type as seen from previous characterization of *Hsf1*/+ growth (Figure 1B, Supplemental Figure 1B and C). *Ts5*/+ as stated in Figure 5C, had increased LER compared to WT. Interestingly, the average LER for the double mutant *Hsf1/+ Ts5*/+ closely matched wild type except for at the 48 hr time point where WT LER slightly exceeded the *Hsf1*/+ *Ts5*/+ LER. (Figure 6A). Analysis of the final leaf lengths of the entire population showed crossing *Hsf1*/+ to *Ts5*/+ reduced final leaf length to wild type lengths (Figure 6B). *Ts5*/+ exhibited a wild type LER and LED growth pattern (Figure 6A).

**Figure 6.**
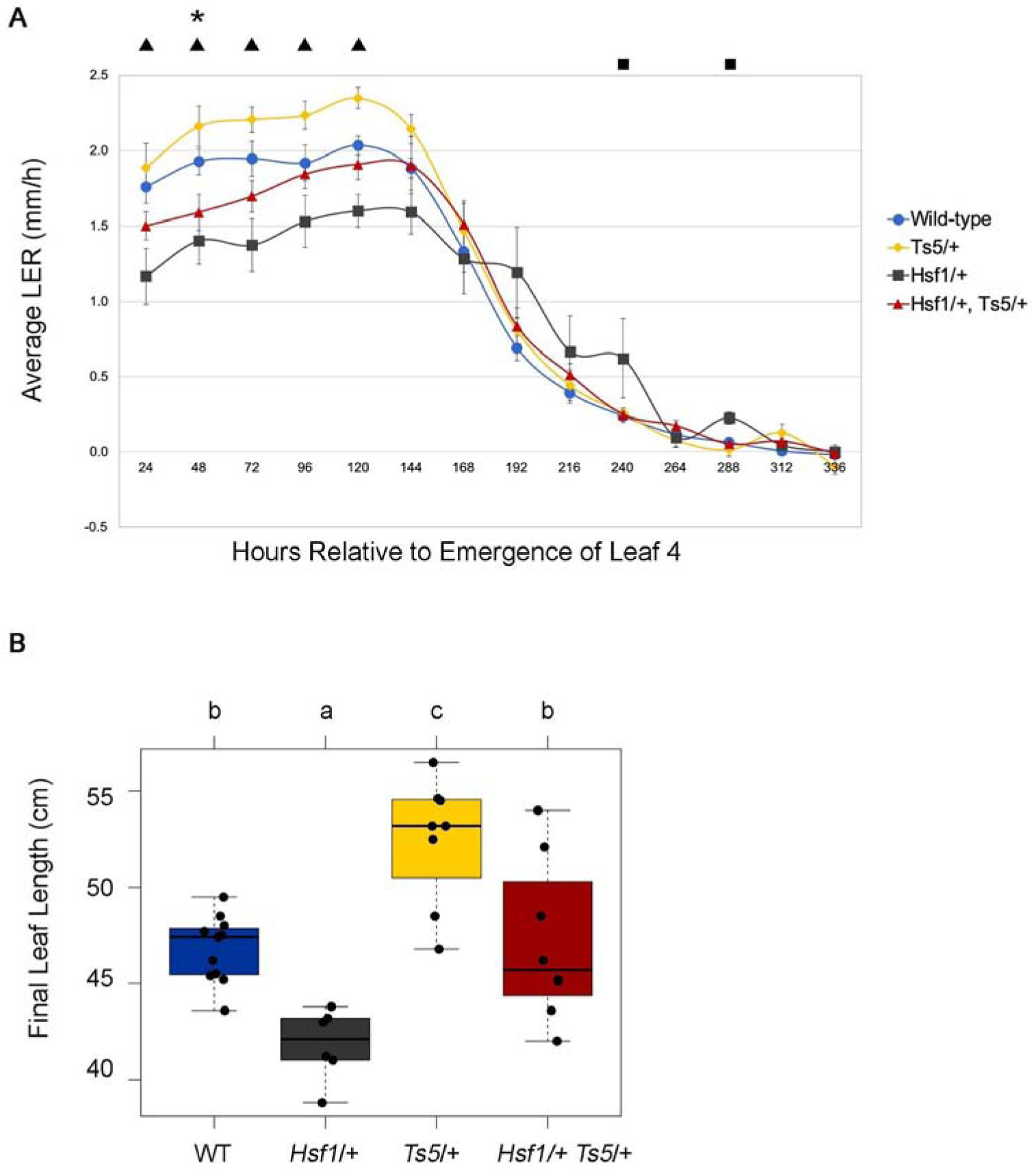
Epistatic interaction of *Hsf1* and *Ts5*. (A) LER of *Hsf1*/+ *Ts5*/+ double mutant compared to WT (asterisk), *Hsf1*/+ (black squares), and *Ts5*/+ (black triangles). Black squares and triangles above the LERs mark significant difference by Student’s t-test p-value ≤ 0.05. Error bars = SE (+/+, n=12; *Hsf1*/+, n=6; *Ts5*/+, n=9, *Hsf1*/+ *Ts5*/+, n=10). (B) Boxplots of sheath length, blade length, and blade width of leaf #1 and #2 of the population described in (A). Horizontal bars represent the maximum, third quantile, median, first quantile, and minimum values respectively. Each dot is a plant.

### Exogenous CK treatment induces expression of JA pathway genes in the leaf growth zone

Since the expression of several JA pathway genes was higher in the leaf GZ of the *Hsf1* CK hypersignaling mutant, we asked if exogenous CK application of maize inbred seedlings could also induce JA pathway expression in the leaf GZ. To do this, 10-day old B73 seedlings were cut at the root:shoot junction, and shoots were incubated for 1, 2 and 4 hours with 10 *µ*M 6-BAP (details in Methods). After incubation, the basal 2 cm of leaf #4, encompassing the DZ and part of the EZ, was collected, and JA pathway expression was quantified using qRT-PCR. We first determined that the exogenous CK application was perceived by assessing expression of three CK early response genes: the type A response regulators *ZmRR3* and *ZmRR6*, and *cytokinin oxidase2* (*ckx2*). Type A response regulators are negative regulators of CK signaling that are rapidly expressed without *de novo* protein synthesis upon CK treatment (To et al., 2004; Ferreira and Kieber, 2005). As expected, *ZmRR6* transcripts were upregulated in the GZ by 1 hour, and all three CK reporters showed robust expression by 4 hours (Figure 7A). Thus, the GZ of leaf #4 was perceiving and responding to the CK application by 4 hours. We next assessed JA pathway expression in these same tissues. Of the genes surveyed, we found an increase in expression of both JA biosynthesis and catabolism genes. Specifically, *ts1, aos1a, aos2a, aoc2, opr7*, and *Ts5* all showed a 1.5 to 3 fold increase in expression after 4 hrs of CK treatment. This showed that CK could induce JA pathway gene expression in the maize leaf GZ after 4 hours.

**Figure 7.**
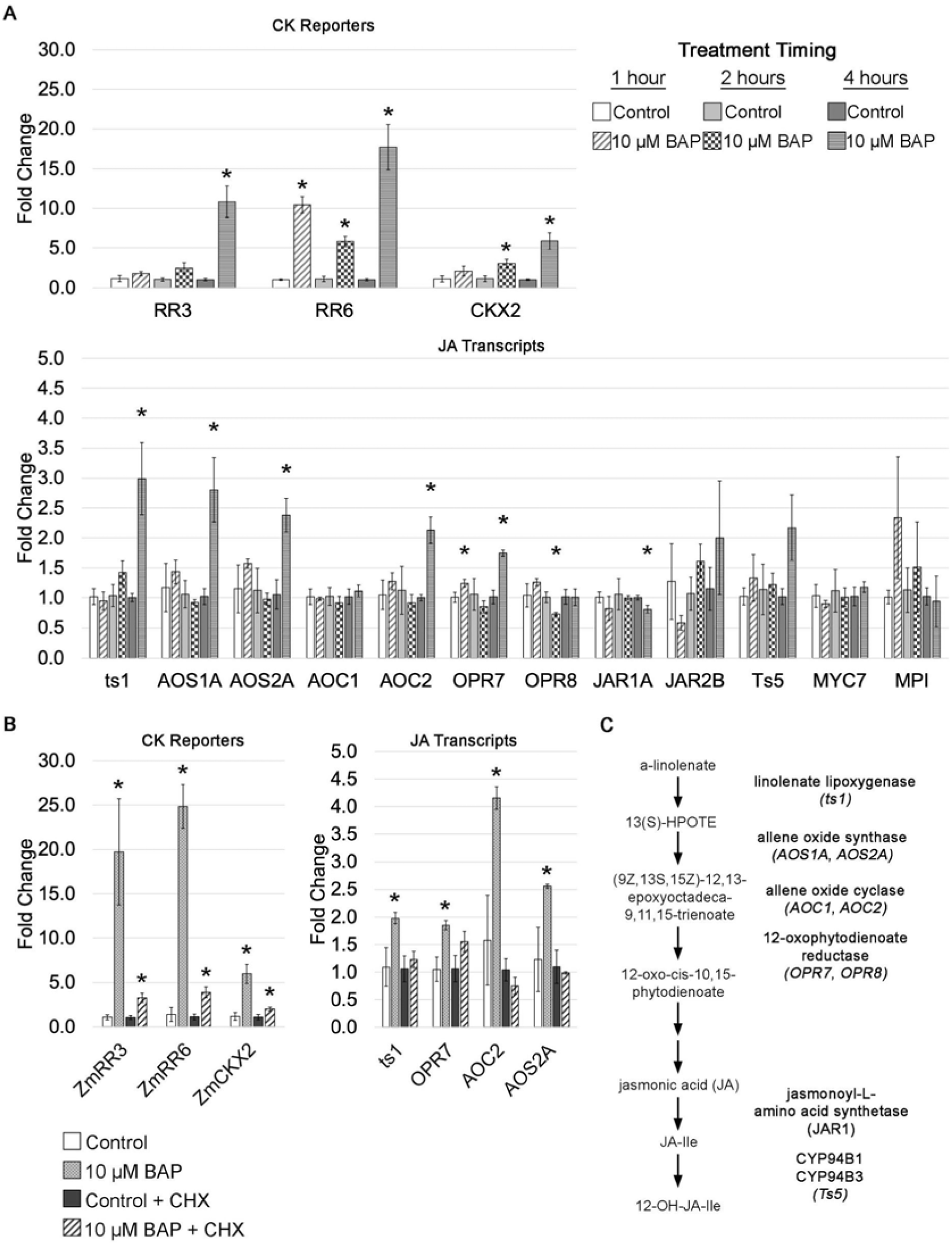
CK induces JA pathway gene expression in the leaf growth zone. **(A)** Quantitative Real-time PCR analysis of CK reporter genes and JA biosynthesis and signaling genes after 10 *µ*M BAP time course. **(B)** Quantitative Real-time PCR analysis of CK reporter genes and JA biosynthesis genes after 10 *µ*M BAP with and without cycloheximide (CHX) treatment. **(C)** Synopsis of JA pathway genes surveyed in (A) and (B). Asterisks in (B) and (C) mark significant difference (P < 0.05) between treatment and respective control.

We next asked if the CK-induced increase in JA gene expression required new protein synthesis downstream of CK signaling. We considered two possibilities: 1) CK treatment and subsequent signaling resulted in the downstream phosphorylation and activation of a transcription factor, such as a type-B response regulator or 2) CK treatment and signaling resulted in the transcription and translation of a new transcription factor that activated expression of the upregulated JA genes. To do this, CK application on cut B73 seedling shoots was repeated with and without cyclohexamide (CHX), a translational blocker. We hypothesized that if CK-induced expression of JA genes was dependent on *de novo* protein synthesis, combined treatment with CK and CHX would result in no increased expression of JA-pathway genes. However, if any JA genes were directly regulated by CK signaling components, like expression of *ZmRR3* and *ZmRR6*, JA gene expression would still be increased in the combined CK and CHX treated samples. We first tested that the combined CK and CHX treatment would work as expected by assessing expression of the three CK reporters. As expected, since type-A response regulator expression does not require *de novo* protein synthesis, *ZmRR3* and *ZmRR6* expression increased in the combined CK and CHX treatment, although the increase was less than with CK alone ol (Figure 6B). In contrast, the CK-induced increased expression of JA pathway genes was abolished with CHX treatment (Figure 6B). This suggests that CK induces transcription and translation of a new protein that regulates JA biosynthesis gene expression in the leaf GZ. Our CK-induction system will be very useful in identifying the CK-induced regulators of these JA pathway genes.

## DISCUSSION

Many dicot examples show that CK signaling promotes the accumulation of plant biomass (Werner et al., 2001; Riefler et al., 2006; Bartrina et al., 2017). *Hsf1*, a monocot, does not follow this pattern and instead shows reduced shoot growth (Muszynski et al., 2019) (Figure 1A and B). In contrast to the constitutive CK receptor mutant in *Arabidopsis*, which has larger leaves with more cells, *Hsf1* has smaller leaves due to a smaller division zone and reduced number of dividing cells (Bartrina et al., 2017) (Figure 1C). Due to the lack of CK signaling mutants in monocots, it is difficult to tell if differential CK-mediated growth responses in monocots and dicots mark true differences in CK signaling or are due to absolute differences in endogenous CK concentrations and perception. However, rice *OsIPT3* transformants overexpressing the rate limiting CK biosynthesis enzyme IPT3, resulted in stunted plants and is another example of excess CK reducing plant growth (Sakamoto et al., 2006).

To understand the connection between CK and leaf growth in *Hsf1*, we focused on characterizing the role of JA in regulating maize leaf growth because of its accumulation in *Hsf1* (Figure 1D and E) and the differential expression of JA biosynthesis genes in the division zone (Figure 2). Previous research has established that monocot and dicot growth is reduced through JA-mediated inhibition of cell proliferation (Yamane et al., 1980; Zhang and Turner, 2008; Noir et al., 2013; Yan et al., 2014). As expected, exogenous application of JA to maize reduced LER which ultimately reduced leaf size (Figure 3A and B). Interestingly, analysis of mutants deficient in JA (*opr7 opr8* and *Ts5*) show increased final leaf size due to increased LER and LED (Yan et al., 2014) (Figure 5). These data show that JA impacts growth primarily by decreasing LER, and support the role of JA mediated growth reduction in *Hsf1* leaves.

Our data suggests that CK hypersignaling induces growth reduction in maize by crosstalk with the growth repressor, JA (Figure 7). Crosstalk between CK and JA is not well characterized and previous data linking the two has been indirect (Ueda and Kato, 1982; Dermastia et al., 1994; O’Brien and Benková, 2013). The majority of these studies relied on exogenous treatments of CK and JA with mixed results that indicated a complex relationship between the two hormones (Ueda and Kato, 1982; Dermastia et al., 1994; O’Brien and Benková, 2013). However, the earliest of these studies observed that JA treatment antagonized CK mediated callus growth (Ueda and Kato, 1982). Our double mutant analysis of *Hsf1*/+ *Ts5*/+ reflect an antagonistic relationship between JA and CK, as the double mutant had wild type LER and final leaf length (Figure 6A and B). In addition, we found that CK treatment of B73 seedlings promotes the transcription and translation of an unidentified protein that promotes the expression of JA biosynthesis genes (Figure 7A and B). Further studies are needed to identify the CK-inducible regulators of the described JA genes.

JA treatments of *Hsf1* suggest that CK crosstalk with other hormones in addition to JA may also play a role in controlling *Hsf1* growth. While crossing the *Ts5*/+ with *Hsf1*/+ rescued the reduced growth phenotype of *Hsf1*/+, the *Hsf1*/+ growth pattern could not be phenocopied with exogenous JA treatment (Figure 4F). These data show that JA treatment reduces wild type leaf size to be equivalent with *Hsf1* (Figure 4A and B) and suggests that JA also reduces leaf size by shortening leaf elongation duration (Figure 4E). Differences between the *Hsf1*/+ *Ts5*/+ cross and the exogenous JA treatment of *Hsf1*/+ may stem from strength of JA perception or reveal the presence of another hormone that crosstalks with CK and JA. Specifically, the extended LED growth pattern is similar to that of a GA signaling mutant, and provides another avenue of hormone crosstalk to investigate in the *Hsf1*/+ mutant (Nelissen et al., 2012). Taken together, it is likely that JA is responsible for reducing LER, and another hormone controls LED in *Hsf1*.

## Conclusion

In conclusion, these data suggest that CK hypersignaling upregulates JA biosynthesis genes, leading to growth reduction in the maize *Hsf1* leaf by suppressing cell proliferation. We provide evidence for an unidentified CK-inducible protein regulator that targets JA biosynthesis genes. Additionally, growth analysis of JA-treated plants and JA-deficient mutants show that JA impacts leaf growth by reducing LER, and removal of JA promotes leaf growth by increasing LER. Collectively, these data highlight a new connection between CK and JA. Determining how CK connects to JA has the potential to provide new insights into the mechanisms plants use to balance growth and defense.

## MATERIALS AND METHODS

### Plant Material, Genetics, Phenotypic Measurements, and Analysis

Inbred B73 was used as the standard maize line for all seed and seedling treatments. The CK hypersignaling mutant *Hsf1-1603* was previously described (Muszynski et al., 2019). The JA-deficient *opr7-5 opr8-2* (we will refer to it as *opr7 opr8*) and *Tasselseed 5 (Ts5)* was previously described in Yan *et al*. 2012 and Lunde *et al*. 2005 respectively (Yan et al., 2012; Lunde et al., 2019). *Hsf1/+* plants were identified by the presence of macrohairs at the V1 stage and prongs in leaf margins past V6 (Muszynski et al., 2019). JA-deficient mutants were grown in flats and genotyped by PCR using the primers described in Supplemental Table S5. Plants were crossed for several generations to produce the following genotypes to analyze: [*+/+, opr7, opr8/+*] WT, [*Hsf1/+, opr7, opr8/+*] CK-hypersignaling only, [+/+, *opr7, opr8*] JA-deficient only, and [*Hsf1/+, opr7, opr8*] CK hypersignaling JA-deficient plants. In parallel, genotypes: [+/+, *ts1/+*] WT, [*Hsf1/+, ts1/+*], CK hypersignaling only, [+/+, *ts1*] JA-deficient, and [*Hsf1/+, ts1*] CK hypersignaling JA-deficient plants were developed. All genotypic classes were grown until leaf 4 matured.

### Standard Germinating Seed Hormone Treatment

A stock and control solution of hormone was made as described by the manufacturer and stored at −80°C. Surface sterilized seeds imbibed overnight were placed embryo-face down, about 20 seeds/petri dish, onto a sterile paper towel and soaked with 2.5 mL of hormone at a working concentration (varied by hormone) in a 15 mm petri dish. Typically, three biological replicates were done per treatment, using 20 seeds/petri dish X 3 = 60 total seeds/treatment. The edges of the petri dishes were sealed with parafilm to prevent evaporation and the entire petri dish was wrapped in foil and placed in a lab drawer for six days. After six days of treatment, germinated seedlings were removed from the petri dish, rinsed with sterile tap water, and transplanted to 1 gallon pots (Sunshine Mix #4 media, supplemented with 2 teaspoons osmocote, 2 teaspoons ironite) and placed in the Pope greenhouse.

### Cytokinin

6-Benzylaminopurine (6-BAP) powder from Sigma Aldrich was first dissolved in 10 drops of 1 N NaOH, and brought to a concentration of 10 mM wither sterile distilled water. A parallel water control stock was also made with 10 drops of 1N NaOH. These stocks were further diluted to achieve the desired hormone treatment concentrations.

### Jasmonic Acid

100 mg of JA (Sigma-Aldrich) was dissolved in 3 mL of 200-proof ethanol and 44.5 mL of sterile ddH_2_O to make a stock concentration of 10 mM JA. A control solution was made by adding 3 mL of 200 proof ethanol to 44.5 mL of ddH_2_O and stored at −80°C. Both the JA and control solutions were diluted with sterile ddH_2_O until the desired working solution concentration was reached. Stock solutions were stored at −80°C in 15 mL tubes. The working solution was made the day treatments started by diluting the 10 mM stock with sterile ddH_2_O to a final volume of 2.5 mL/petri dish.

### Final Leaf Size Measurements

Treated seedlings were grown until the fifth leaf was completely collared (the auricle and ligule that defines the junction between the leaf sheath and blade was visible), ensuring that leaves #1 to #4 had completed growth. Sheath length, blade length, and blade width were measured for leaves #1 (most basal, first formed) to leaf #4. Leaves were measured by harvesting each leaf at its insertion into the stem. For sheath length— length was measured from the base of the sheath to the point at which the sheath transitions to the auricle at the midline of the leaf. For blade length— length was measured along the midrib from the auricle to the distal blade tip. For blade width— width was measured at the midpoint of blade length across the blade from margin to margin.

### Growth Rate Measurement

Leaf elongation rates (LER) were taken when leaf #4 emerged from the whorl and was at steady-state growth, when LER is constant (Sun et al., 2017). Briefly, the length of leaf #4 was measured as the distance from the insertion point of leaf #1 at the base of the plant to the tip of leaf #4 every 12 or 24 hours until leaf #4 stopped growth (leaf length did not change for 2-3 consecutive time points). LER was calculated by dividing the difference in leaf length (cm) by the time elapsed (24 hrs). Leaf elongation duration (LED), the measure of time from when the leaf is 10 cm to final length, was determined from plotting LER by time elapsed. Leaf elongation duration (LED) was determined when steady state growth stopped as observed when plotting LER by days post leaf 4 emergence from the whorl. Finally, plants were dissected and leaf blade length, leaf blade width (measure at ½ the blade length mark), and leaf sheath length were measured on leaves #1 – 4.

### Seedling treatments and JA-pathway gene expression analysis

Seedling treatments were performed as described in (Giulini et al., 2004) on B73 seedlings when leaf 4 was emerging from the whorl. Briefly, individual seedlings were cut at the shoot-root junction and submerged in 500 uL of 10 uM 6-BAP or equivalent control for 4 hrs. The basal 2 cm of the leaf, were division and expansion occurs, was dissected and put in 500 ul of IBI Isolate (IBI Scientific, CAT: IB47601) for RNA extraction following the manufacturer’s recommendations. RNA was quantified by using ND-1000 Spectrophotometer (Nanodrop, Wilmington, DE). A total of 2 ug of RNA was used to synthesize cDNA with SuperScript IV VILO Master Mix with ezDNase Enzyme kit (Thermo Fisher Scientific, CAT: 11766050) following the manufacturer’s recommendations. Finally, 1:10 dilution of cDNA was used for RT- and quantitative RT-PCR.

Samples were initially screened for CK perception by RT-PCR amplifying *ZmRR3* (*abph1*; Zm00001d002982), a type-A response regulator that is only expressed when CK is present (Giulini et al., 2004), using the EconoTaq® PLUS GREEN 2X Master Mix (Lucigen; Middleton, WI) and following the manufacturer’s recommendations. The RT-PCR was performed using S1000™ Thermal Cyclers (Bio-Rad; Hercules, CA) using the following cycling program: step 1 = 98 °C for 2 min, step 2 = 98 °C for 30 sec, step 3 = 60 °C for 30 sec, step 4 = 72 °C for 30 sec, step 5 = repeat steps 2 – 4 29 times, step 6 = 72 °C for 5 min, and step 7 = 10 °C. PCR products were run in 2% agarose gel electrophoresis using a 100 bp DNA ladder (GenScript; CAT: M102O).

Once perception was confirmed, genes that encode for the biosynthetic enzymes along the JA-pathway were evaluated by quantitative RT-PCR using the iQ SYBR Green Supermix (Bio-Rad; CAT: 1708882) reagents, following manufacturer recommendations, and Bio-Rad CFX96 Touch™ thermocycler (Bio-Rad; Hercules, CA) with primers listed in Supplemental Table 5. Cq values were used to calculate Fold Change differences between the control and the treatments following (Livak and Schmittgen, 2001) and calculating significant differences using Student’s t-test.

### Hormone Analysis

#### Plant metabolite assays

Plant hormones (cytokinins, jasmonate, salicylic acid, auxin, *cis*-zeatin, *trans*-zeatin) were measured by HPLC-mass spectrometry (HPLC-MS) as described previously (Schäfer et al., 2016).

## ACKNOWLEDGEMENTS

We would like to thank all past and present members of the Muszynski Lab for their help in data collection including Dylan Oates, Miranda Yip, Bridnie Hill, Jessica Szyska and Sirut Buasai. ANU was awarded a Syngenta Agricultural Scholarship.

Hormone profiling done by GJ was funded by US National Science Foundation award 1339237 and a Friedrich Wilhelm Bessel Research Award from the Humboldt Foundation.

## SUPPLEMENTAL DATA

**Supplementary Figure S1.**
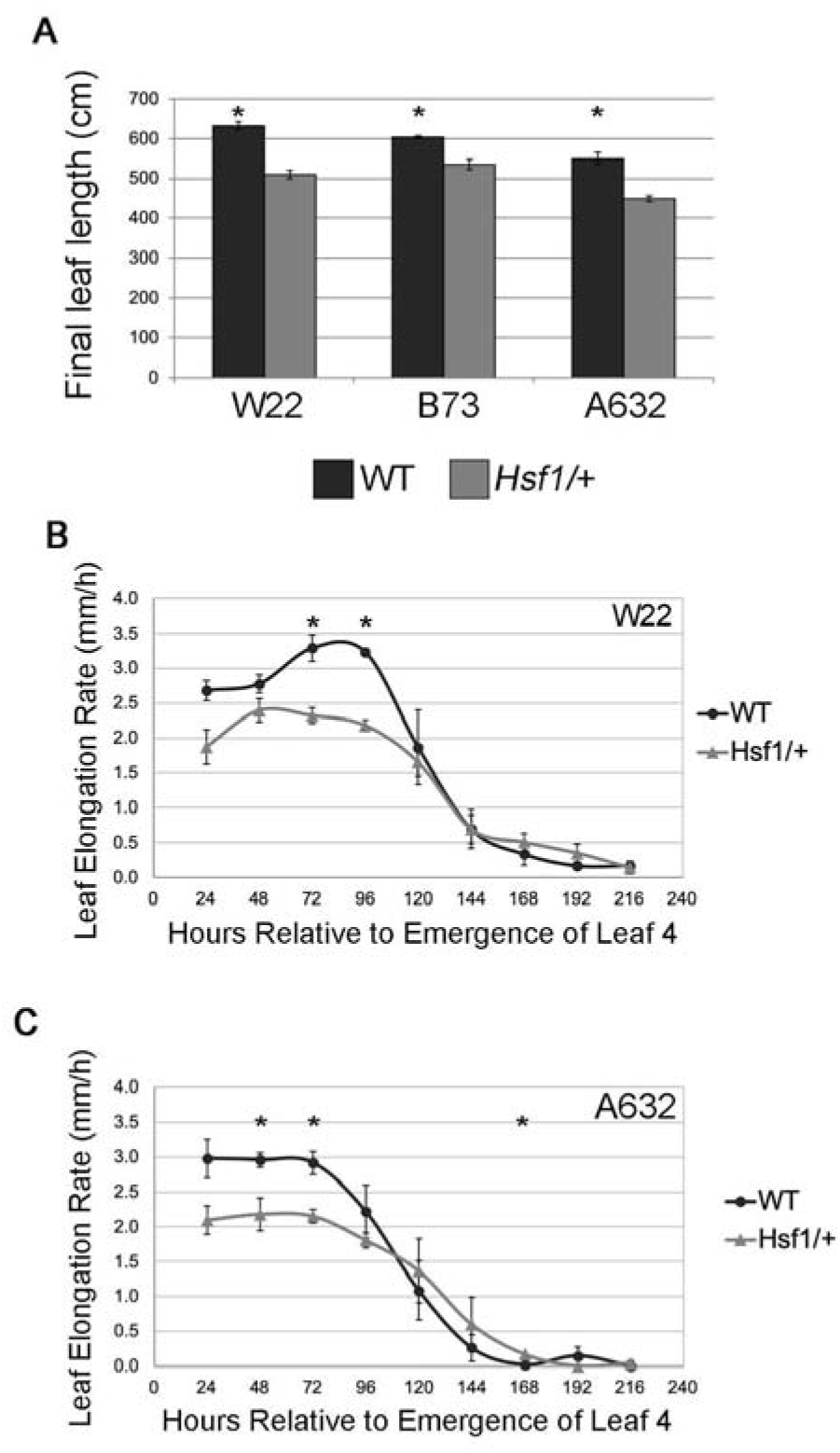
*Hsf1* growth in different inbred backgrounds. **(A)** Barplots of WT and *Hsf1*/+ final leaf lengths. Error bars = SE. **(B-C)** Average leaf elongation rate (LER) of leaf #4 of *Hsf1*/+ and WT-siblings in the **(B)** W22, and **(C)** A632 inbred backgrounds. Asterisks mark significant difference P < 0.05. Error bars = SE.

**Supplemental Table S1.**
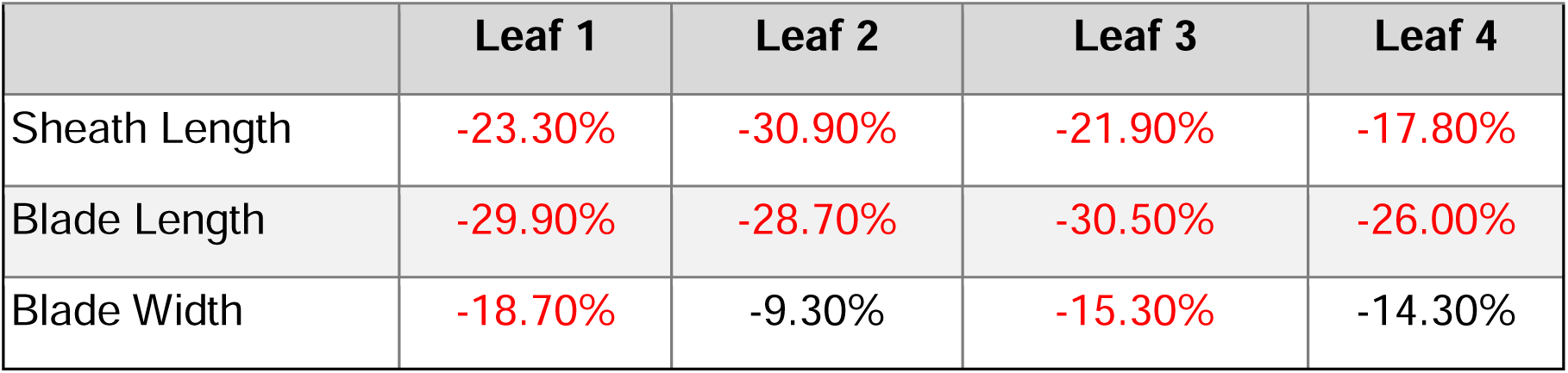
Percent leaf size reduction after exogenous 1 mM JA treatment. Percent reductions [(JA-C)/C *100] in sheath length, blade length, and blade width by leaf number. Red means significant value P < 0.05.

|  | Leaf 1 | Leaf 2 | Leaf 3 | Leaf 4 |
| --- | --- | --- | --- | --- |
| Sheath Length | -23.30% | -30.90% | -21.90% | -17.80% |
| Blade Length | -29.90% | -28.70% | -30.50% | -26.00% |
| Blade Width | -18.70% | -9.30% | -15.30% | -14.30% |

**Supplementary Figure S2.**
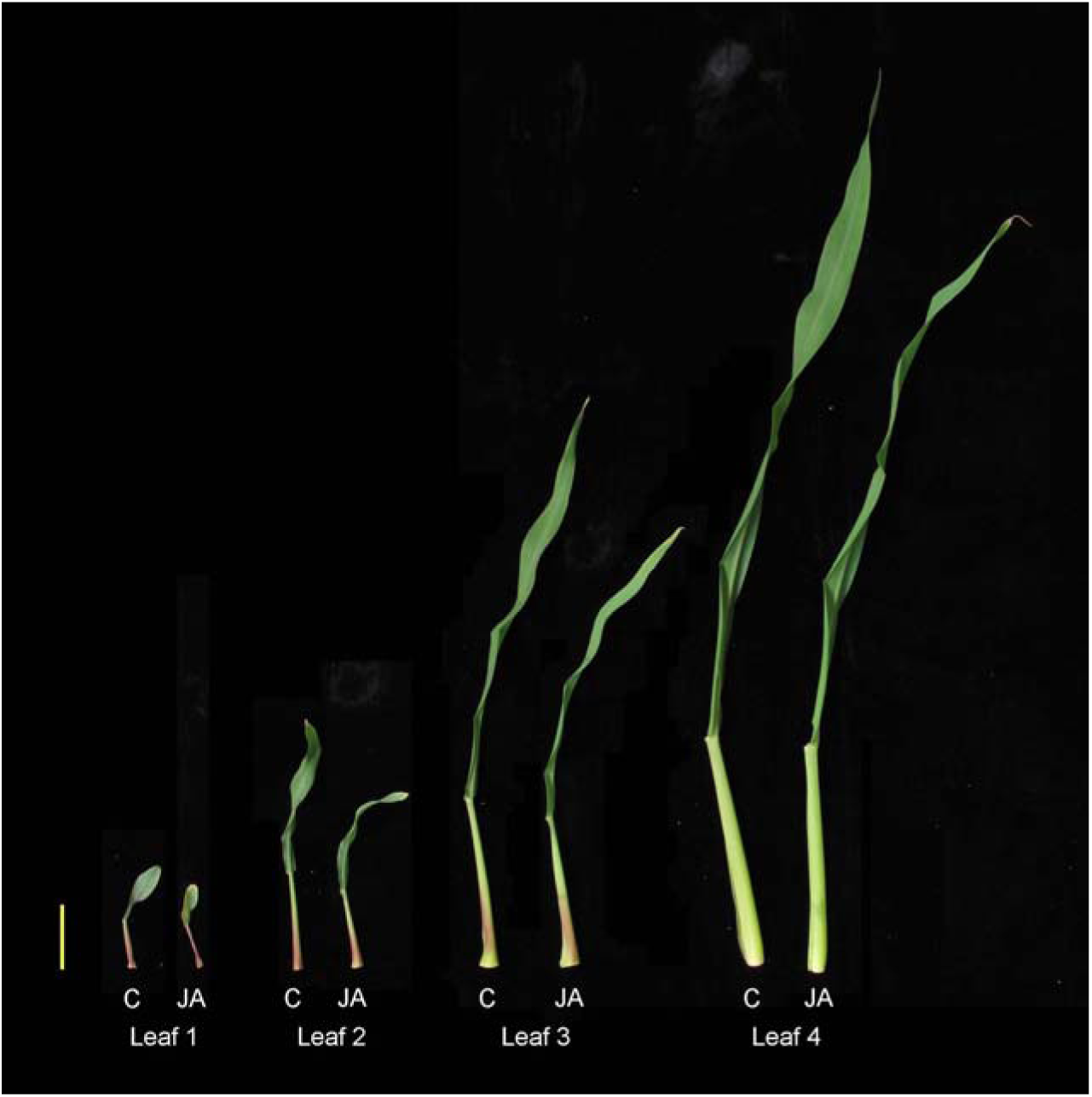
Comparison of control (C) and jasmonic acid (JA) treated leaves #1-4. Scale bar = 5 cm.

**Supplemental Figure S3.**
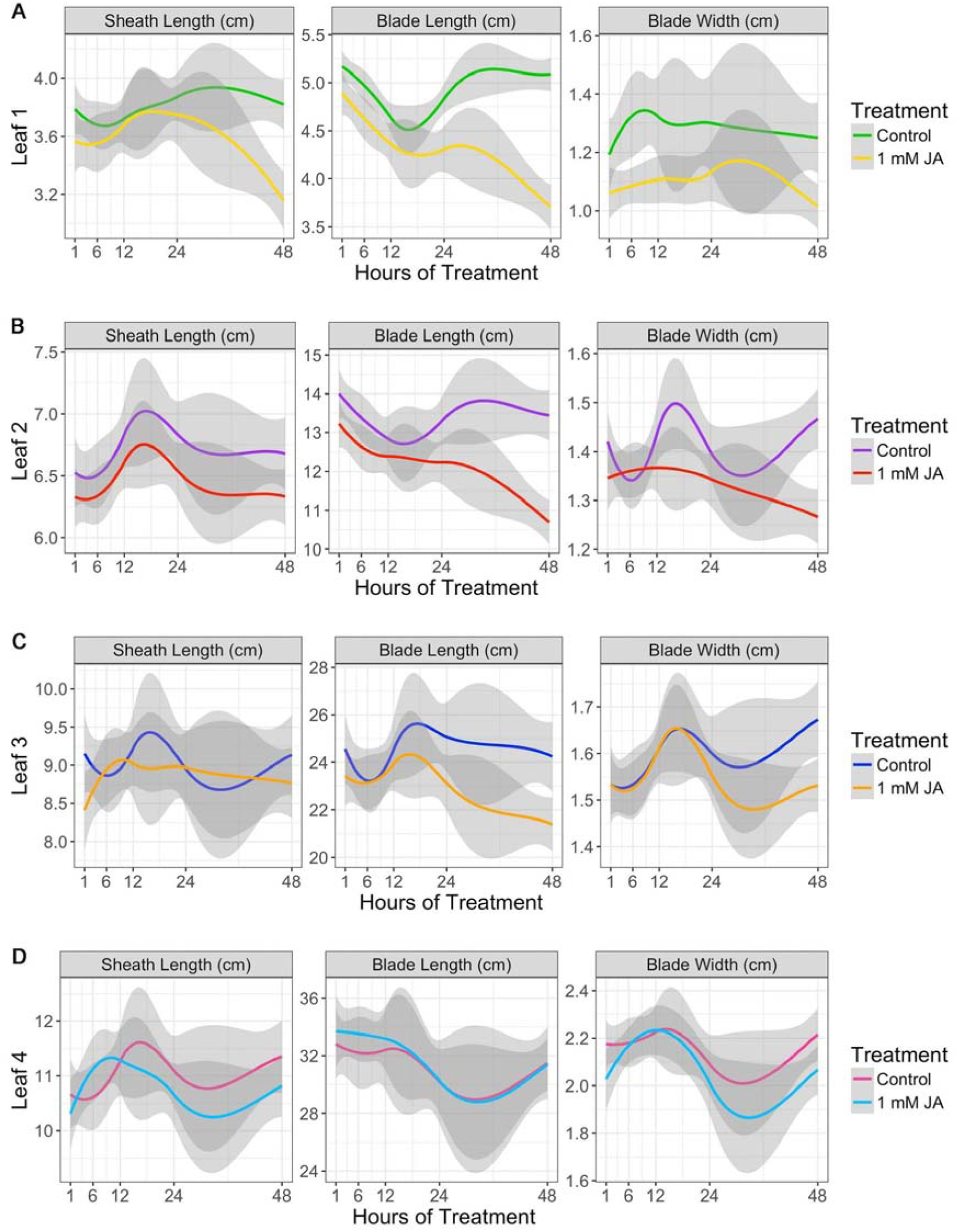
Final leaf measurements of B73 treated with 1 mM JA or control solution for 1, 6, 12, 24, or 48 hours. Leaf 1 (A), leaf 2 (B), leaf 3 (C), and leaf 4 (D) were measured for all plants. Each dot is plant, lines are smoothed conditional means, and shaded area is the 95% confidence interval. Treatments are significant where confidence intervals do not overlap.

**Supplemental Table S2.** Percent leaf size reduction after 48 hours of exogenous 1 mM JA treatment. Percent reductions [(JA-C)/C *100] in sheath length, blade length, and blade width by leaf number. Red means significant value P < 0.05.

|  | Leaf 1 | Leaf 2 | Leaf 3 | Leaf 4 |
| --- | --- | --- | --- | --- |
| Sheath Length | -17.30% | -5.20% | -4.00% | -4.70% |
| Blade Length | -27.10% | -20.50% | -11.70% | -0.30% |
| Blade Width | -18.80% | -13.60% | -8.50% | -6.70% |

**Supplemental Figure S4.**
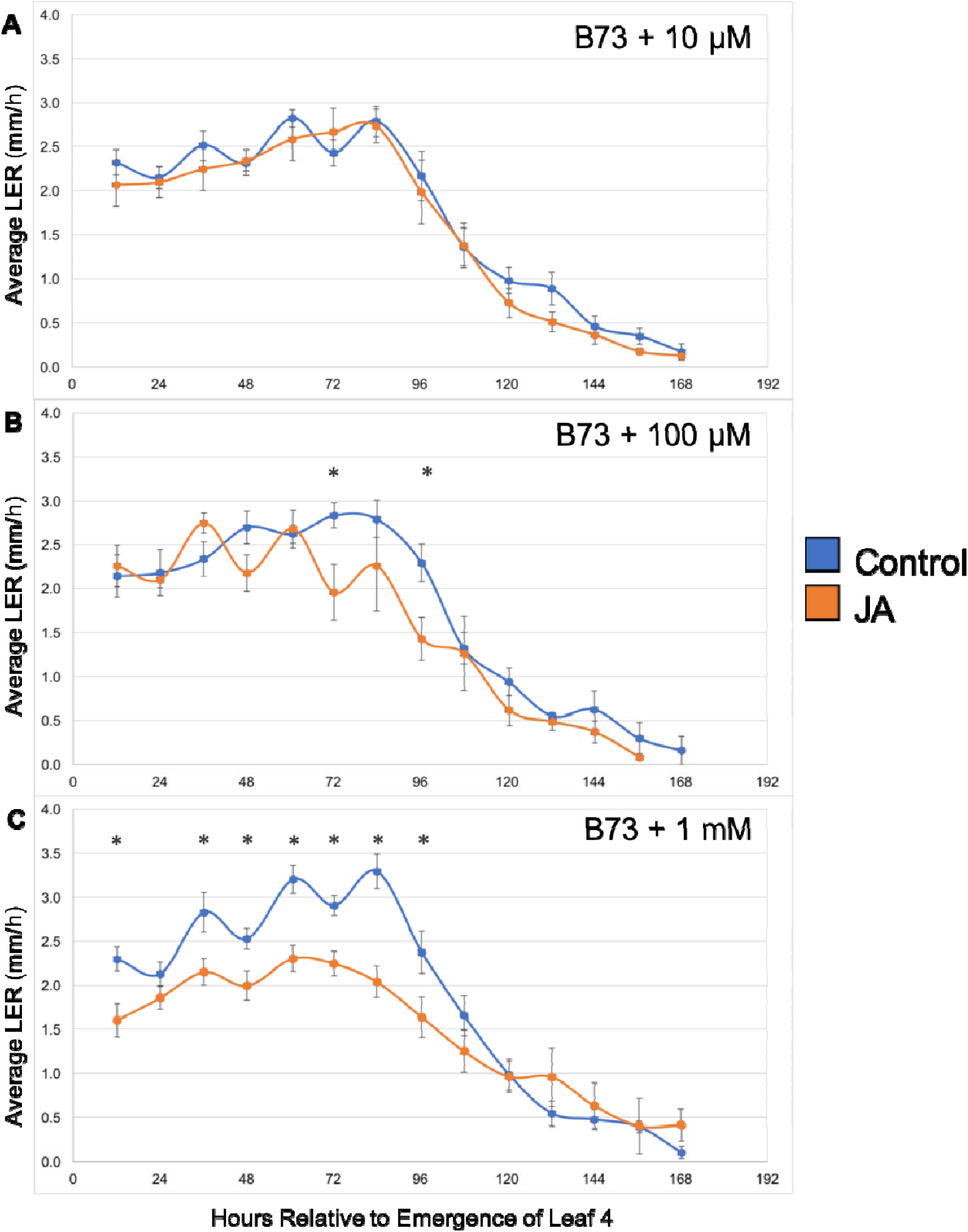
LER dose response to JA in B73. Leaf 4 LERs of B73 treated with (A) 10 *µ*M, (B) 100 *µ*M, and (C) 1 mM JA for 6 days. Significant differences by Student’s t-test are marked by asterisks and error bars are SE.

**Supplemental Table S3.** Relevant comparisons of *Hsf1*/+ and WT-sibling final leaf size percent reductions after JA treatment. (i) WT-sibling compared to *Hsf1/+* without JA, (ii) WT-sibling with JA treatment, (iii) *Hsf1/+* with JA treatment, (iv) WT-sibling compared to *Hsf1/+* both treated with JA. Red means significant percent difference P < 0.05.

| **i. WT Control vs. *Hsf1/+* Control** |  |  |  |  |
| --- | --- | --- | --- | --- |
|  | **Leaf 1** | **Leaf 2** | **Leaf 3** | **Leaf 4** |
| Sheath Length | -22.20% | -18.40% | -16.50% | -14.50% |
| Blade Length | -20.20% | -21.90% | -22.80% | -13.50% |
| Blade Width | -22.90% | -20.60% | -21.80% | -15.20% |
| **ii. WT Control vs. WT JA** |  |  |  |  |
|  | **Leaf 1** | **Leaf 2** | **Leaf 3** | **Leaf 4** |
| Sheath Length | -28.80% | -16.90% | -15.90% | -12.20% |
| Blade Length | -44.80% | -39.00% | -25.80% | -18.90% |
| Blade Width | -27.80% | -23.40% | -20.40% | -10.60% |
| **iii. *Hsf1/+* Control vs. *Hsf1/+* JA** |  |  |  |  |
|  | **Leaf 1** | **Leaf 2** | **Leaf 3** | **Leaf 4** |
| Sheath Length | -16.50% | -15.80% | -15.00% | -7.80% |
| Blade Length | -38.10% | -37.40% | -25.30% | -17.50% |
| Blade Width | -29.00% | -9.20% | -17.60% | -13.40% |
| **iv. WT JA vs. *Hsf1/+* JA** |  |  |  |  |
|  | **Leaf 1** | **Leaf 2** | **Leaf 3** | **Leaf 4** |
| Sheath Length | -8.80% | -17.30% | -15.60% | -10.20% |
| Blade Length | -10.50% | -19.80% | -22.40% | -12.00% |
| Blade Width | -24.20% | -5.90% | -19.10% | -17.90% |

**Supplemental Table S4.** Relevant comparisons of *opr7 opr8* double mutant final leaf size percent reductions. (i) *opr7 opr8* compared to JA sufficient *opr7/opr7 OPR8/opr8* (ii) *opr7 opr8* compared to JA sufficient *opr7/opr7 OPR8/OPR8*. Red means significant percent difference P < 0.05.

| **i. *opr7/opr7 opr8/opr8* vs. *opr7/opr7 OPR8/opr8*** |  |  |  |  |
| --- | --- | --- | --- | --- |
|  | **Leaf 1** | **Leaf 2** | **Leaf 3** | **Leaf 4** |
| Sheath Length | 39.5% | 22.2% | 11.0% | -1.4% |
| Blade Length | 43.0% | 36.8% | 22.6% | 14.2% |
| Blade Width | -7.6% | -1.6% | 10.0% | 17.6% |
| **ii. *opr7/opr7 opr8/opr8* vs. *opr7/opr7 OPR8/OPR8*** |  |  |  |  |
|  | **Leaf 1** | **Leaf 2** | **Leaf 3** | **Leaf 4** |
| Sheath Length | 43.3% | 21.0% | 12.4% | -2.3% |
| Blade Length | 48.2% | 38.2% | 24.2% | 12.9% |
| Blade Width | -9.5% | -4.8% | 9.2% | 18.6% |

**Supplemental Table S5.**
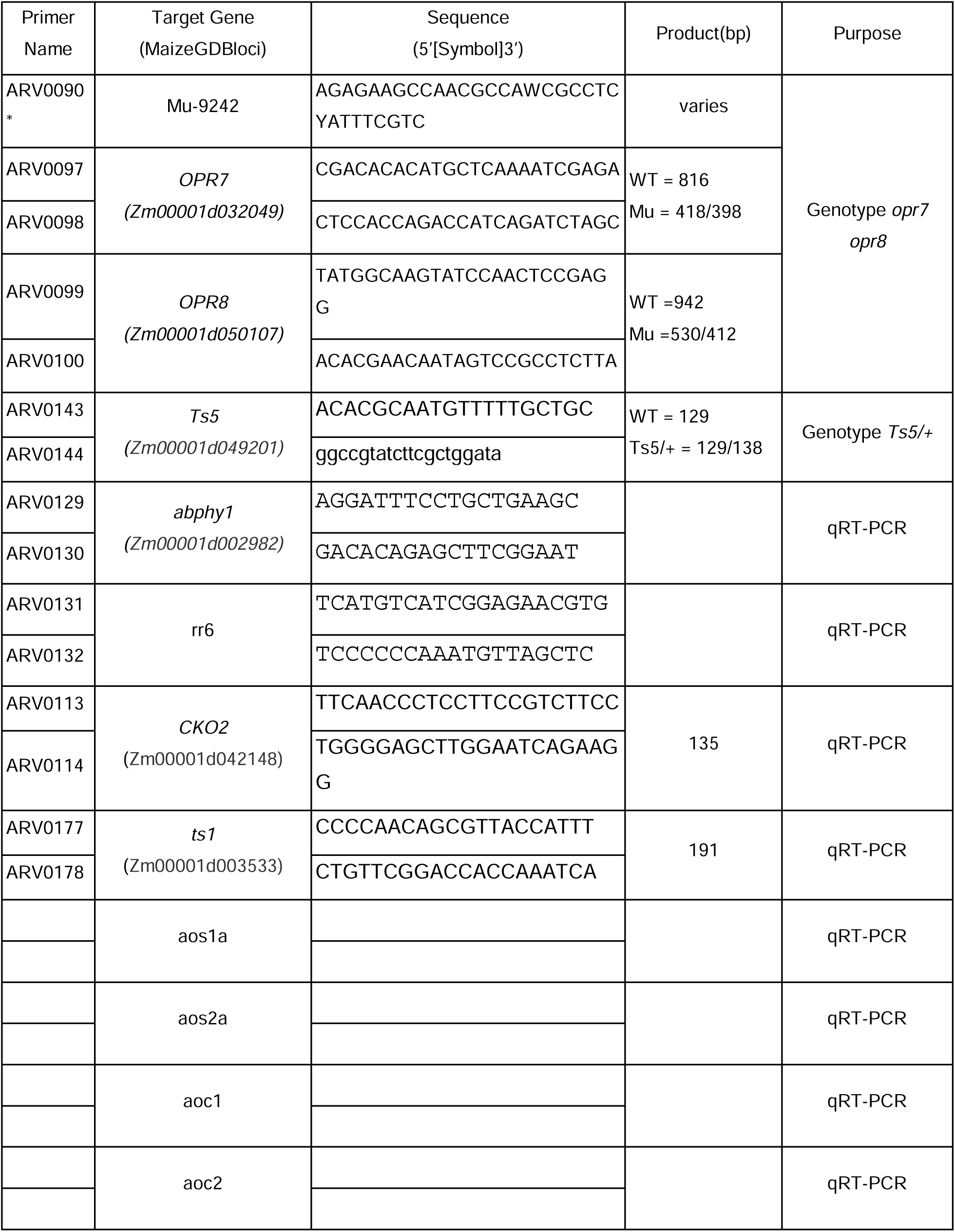
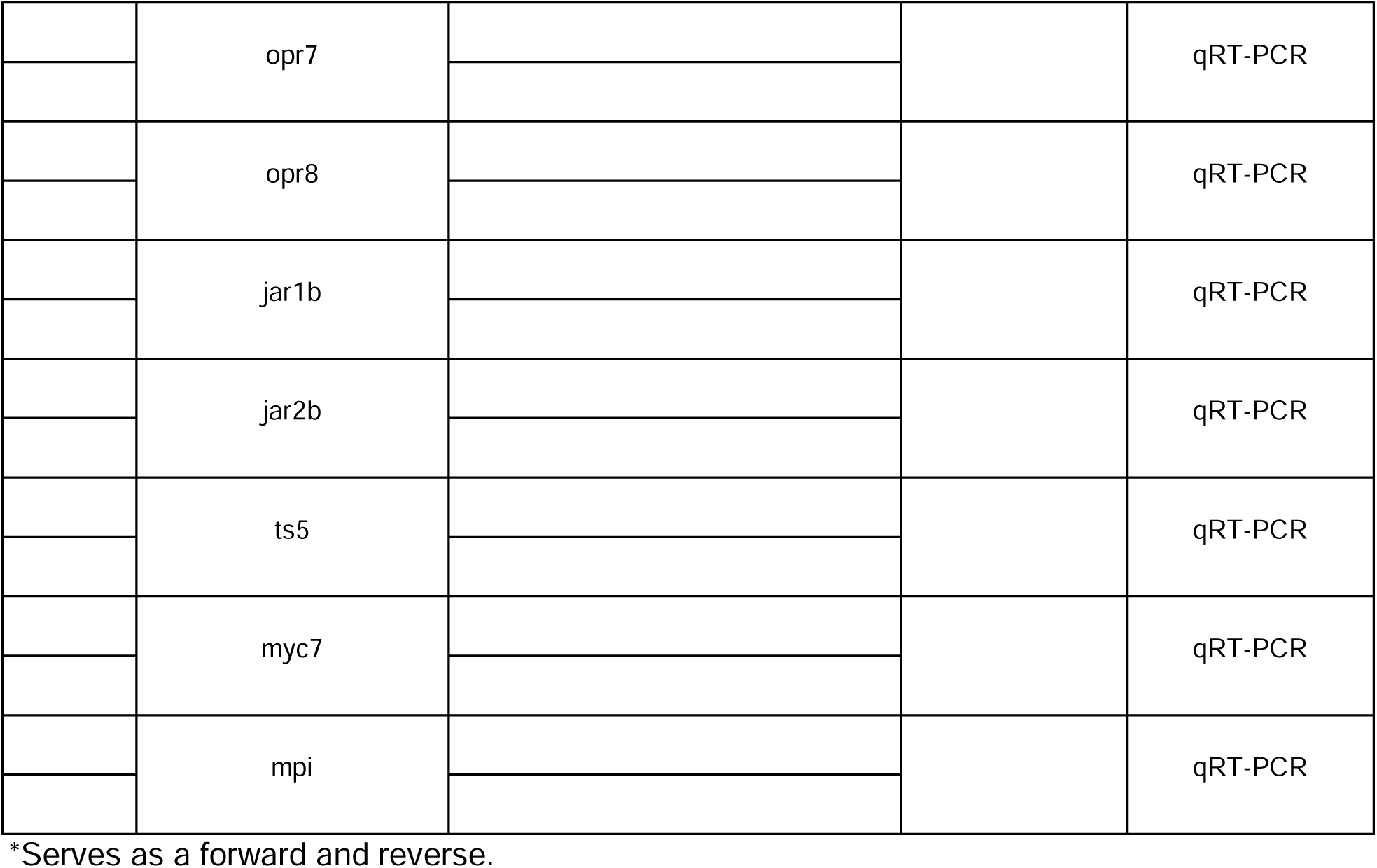
The list of primers used in this study.

